# A comparison of asymmetric before-after control impact (aBACI) and staircase experimental designs for testing the effectiveness of stream restoration

**DOI:** 10.1101/359406

**Authors:** Tom M. Loughin, Stephen N. Bennett, Nicolaas W. Bouwes

## Abstract

Before-after-control-impact (BACI) experimental designs are commonly used in large-scale experiments to test for environmental impacts. However, high natural variability of environmental conditions and populations, and low replication in both treatment and control areas in time and space hampers detection of responses. We compare the power of two asymmetric BACI (aBACI) designs to two staircase designs for detecting changes in juvenile steelhead (*Oncorhynchus mykiss*) abundance associated with a watershed-scale stream restoration experiment. We performed a simulation study to estimate the effect of a 25% increase in steelhead abundance using spatial and temporal estimates of variance from an ongoing study, and determined the power of each design. Experimental designs were then applied to three streams and each stream was composed of three 4 km long *sections*. We compared the power of a single treatment section in one stream (BACI-1), three simultaneous treatments of all sections in one stream (BACI-3), three sequential treatments in one stream (STAIRCASE-1), and three sequential treatments in one section in each stream (STAIRCASE-3). All designs had ≥ 94% power to detect a 25% increase in abundance assuming average variance. Under worst-case variance (i.e., upper 95% confidence limits of historical variance estimates), the STAIRCASE-3 design outperformed the BACI-1, BACI-3, and STAIRCASE-1 designs (i.e., 77%, 41%, 8%, and 33% power respectively). All the designs estimated the effect of the simulated 25% abundance increase, but the length of the confidence interval was much shorter for the STAIRCASE-3 design compared to the other designs, which had confidence intervals 58-596% longer. The STAIRCASE-3 design continued to have high power (88%) to detect a 10% change in abundance, but the power of the other designs was much lower (range 34-56%). Our study demonstrates that staircase designs can have significant advantages over BACI designs and therefore should be more widely used for testing environmental impacts.

## Introduction

Impacts from manipulations to ecosystems, such as the extraction of natural resources, must be identified and quantified to develop strategies to reduce our footprint on the environment and manage resources sustainably. Ecosystem experiments using impacts (e.g., perturbations such as logging, addition of nutrients) have led to a greater understanding of the influence of management actions on ecosystem processes and biological populations [1–3]. Ecosystem experiments have provided a wealth of information because they were appropriately scaled – the impacts were large (e.g., often whole watersheds) and the monitoring was intensive (e.g., monitoring multiple scales and for many years or decades), which allowed detection of environmental changes that would not have been possible had the studies been conducted on smaller spatial and temporal scales [4, 5]. However, ecosystem scale experiments are expensive, difficult to replicate, and require large changes (impacts) to reliably detect a response. Extrapolation of results from ecosystem experiments is also challenging because of the lack of replication, unless mechanisms of change are determined (typically through directed studies). Despite these challenges, ecosystem style experiments have recently been advocated for determining the effectiveness of restoration actions because of the difficulty in determining how habitat and populations change due to natural variability in space and time [6].

Attempts to restore degraded habitats is exemplified in the Pacific Northwest, where hundreds of millions of dollars have been spent restoring stream habitat in an effort to increase the production of salmon and steelhead [7]. However, much of the restoration and its associated monitoring have been implemented at small scales (e.g., sites < 1000 m), and scattered across multiple watersheds, which has made it difficult to determine if site scale responses can be scaled to larger areas [e.g., increases in production of an entire stream; 8, 9, 10]. Thus, implementation of experiments that apply restoration treatments across a large portion of a watershed, coupled with watershed wide monitoring to increase our understanding of the effectiveness of these strategies is gaining popularity [11, 12].

A common experimental design to improve our ability to detect the response of an impact or restoration treatment is to use a before-after-control-impact (BACI) design, in which a treatment unit is compared to a control or reference unit before and after the treatment is applied [13]. BACI designs have been used to detect large and permanent changes in the mean of a population resulting from an impact [14]. However, BACI designs perform poorly or provide misleading results when the potential changes after an impact are small or gradual (i.e. not a step change), when the behavior of the control system is quite different, or when the changes are not to the mean but to the variability of the population [15, 16]. As with any experiment, BACI designs become more powerful when more replicates are used; sometimes referred to as Multiple BACI or mBACI (Downs et al 2002). Underwood (1994) argues that asymmetric BACI (aBACI) designs, where a treatment system is compared to multiple controls is much more powerful than the original BACI design but does not require multiple treatments. Incorporating multiple control sites also protects the experiment from the loss of a single control site or having a poor control site that doesn’t compare well to the treatment site regardless of the treatment [17, 18].

Temporal replication of the impact is also required if the results of an experiment are to be extrapolated beyond specific initial conditions, and the time sequence in which observations were taken. Environmental factors such as extreme weather events can affect the magnitude of an impact and subsequent effect on the response metric (e.g., population abundance). Thus, the effect of an impact cannot be distinguished from a random time x impact interaction where the impact was established during a single time period [19, 20]. However, temporal replication of long-term mBACI experiments (i.e. several mBACI experiments, each with a different start date) may be financially and logistically prohibitive.

An alternative to the mBACI design is the “staircase” design, first proposed by Walters et al. (1988), which involves a modification to the BACI design whereby impacts are staggered in time in different replicates. In this way, each replicate experiences a different sequence of random environmental effects in the periods before and after the impact. This allows for replication of impacts and estimation of the resulting effect through space and time with less effort than mBACI replicated through time.

Numerous large-scale, long-term restoration experiments, referred to as Intensively Monitored Watershed (IMW) studies, have been initiated in the Pacific Northwest in response to a lack of clear evidence that millions of dollars in stream restoration has had a positive effect of populations of Endangered Species Act listed salmon and steelhead [https://www.pnamp.org/project/3133; 6, 7, 21]. The aBACI experimental designs, often with one treatment watershed and one or more control watersheds (i.e., asymmetrical BACI), are the dominant designs that are being used to evaluate these stream restoration experiments.

We present a comparison of the statistical power of two aBACI and two staircase designs to detect the effectiveness of stream restoration at increasing the abundance of juvenile steelhead (*Oncorhynchus mykiss*). We assess statistical power by simulating abundance measurements with properties that are similar to those observed in an IMW we are currently implementing. We assign “true” treatment effects to these measurements, artificially increasing the abundances according to when and where the treatments would be applied in experiments following the candidate designs. We then use design-appropriate ANOVA models on the measurements. We repeat this process many times and estimate the design’s statistical power from the proportion of repetitions in which the treatment effect is detected by the analysis.

Our goal is to use a simulation models to test the power of different experimental designs and provide researchers with alternative approaches to conducting experiments that can maximize learning while trying to achieve the goal of species recovery. In particular, our objectives were to determine the statistical power of aBACI and staircase designs to detect a range of changes (5-40% increase) in juvenile steelhead abundance within a 12 year restoration experiment. We developed four different experimental designs that could be generalized to other studies: 1) aBACI single treatment section in one stream with 8 control sections (BACI-1); 2) aBACI three simultaneous treatment sections in one stream and 6 control sections (BACI-3); 3) single stream staircase with three sequential treatments in one stream with 6 control sections (STAIRCASE-1); and 4) three stream staircase with three sequential treatments one in each of three streams with 6 control sections (STAIRCASE-3). Results from these simulations will help managers develop robust experimental designs that have a greater probability of detecting restoration effects.

## Methods

We performed a simulation study to determine the statistical power of four experimental designs to detect a known juvenile steelhead response to restoration. The simulation approach included: 1) estimating spatial and temporal variance for juvenile steelhead abundance in an ongoing IMW to create confidence intervals for each variance component, 2) building a simulation model of the general IMW layout based on the study setting and specified structure of blocking factors and responses, 3) using a Monte-Carlo simulation to generate data of fish abundance responses at each possible sampling time and location, using three levels of variance, 4) adding multiple systematic fixed treatment effects to treated sections of each model watershed according to each of the experimental designs, and 5) analyzing the results using the appropriate model for each design and recording whether each treatment effect was detected in each analyses.

### Asotin Creek Data Collection and Compilation

We used the Asotin Creek Intensively Monitored Watershed as a case study to assess the performance of the aBACI and staircase experimental designs. Asotin Creek is a tributary to the Snake River in southeast Washington in the Columbia Plateau and Blue Mountains ecoregion (Fig 1). The Asotin Creek IMW study area consists of three sub-watersheds: Charley, North Fork Asotin, and South Fork Asotin Creeks. The addition of large woody debris (LWD) was proposed as the main restoration treatment to increase habitat complexity, which was then expected to increase juvenile steelhead abundance [22, 23].

**Fig 1.**
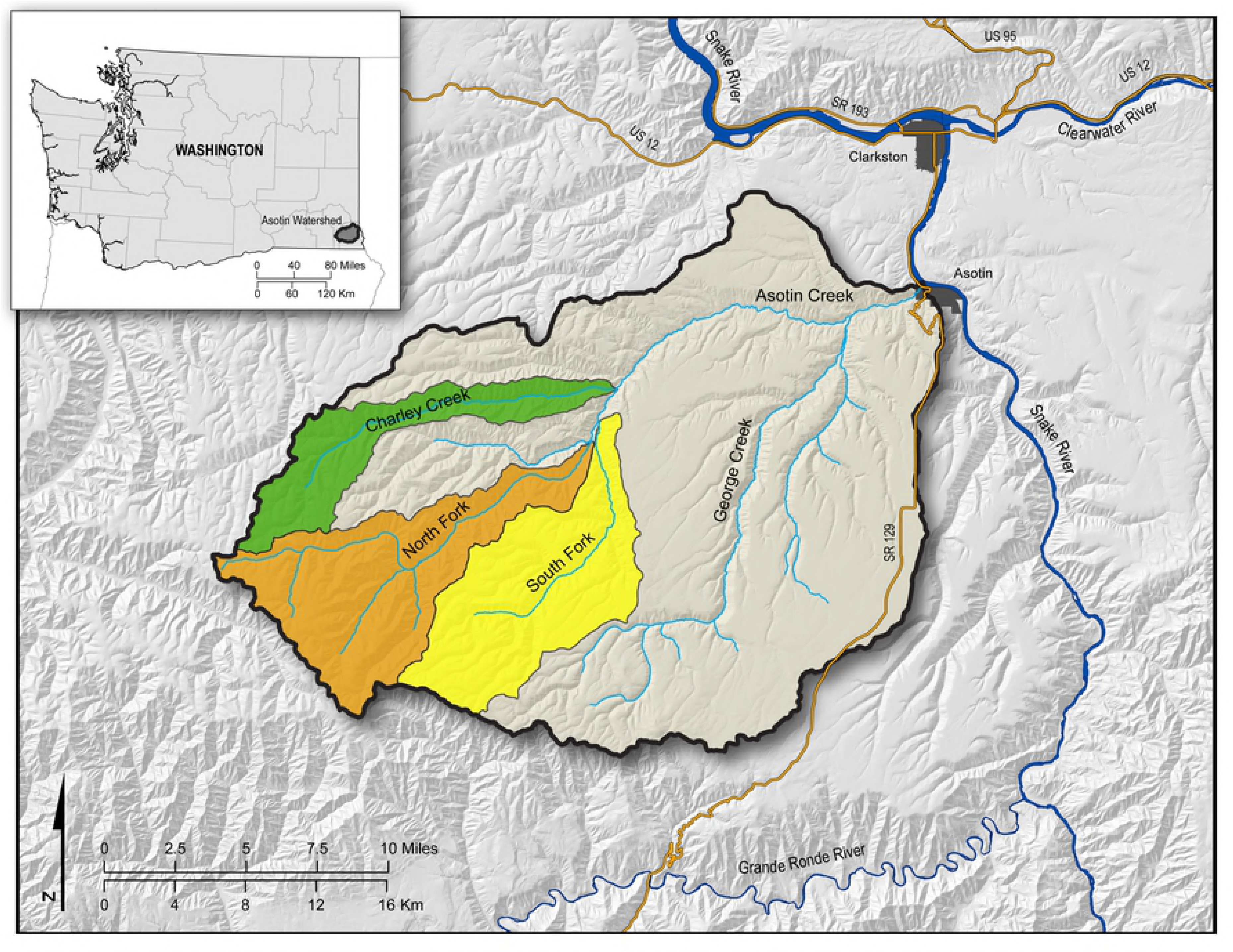
Location of Asotin Creek within Washington and the Asotin Creek Intensively Monitored Watershed study watersheds (highlighted in color) within the Asotin Creek.

The lower 12 km of each study stream was chosen as the IMW study area because 1) the majority of steelhead tributary adult spawning and juvenile rearing takes place in Asotin Creek; 2) access to the upper watershed is difficult and would limit monitoring abilities; and 3) the stream channel and valley conditions (i.e., reach types) in the lower 12 km of each study stream is relatively homogeneous. The size of the sections (experimental units) were based on 1) the size of the treatment area that we believed was a reasonable length of stream that could be restored within a year and 2) would likely ensure that both the experimental “populations” and the stream effects from the restoration were independent between sections, and 3) was likely to capture the mechanisms that would affect the population of interest (i.e., the changes in fish habitat that would lead to changes in fish abundance). We estimated that approximately 150-200 LWD structures could be built per 4 km treatment section to increase wood frequency to historic levels [24, 25]. We used monitoring information on juvenile home ranges to confirm that relative to the population size within the sections, the exchange rate of individuals (fish ≥ 70 mm tagged with passive integrated transponders) between sections was low, and therefore we considered the fish response independent across all sections. We also expected that the hypothesized geomorphic response would be relatively localized [26]. Based on these criteria, each study stream was divided into three 4-km long sections for a total of nine sections (Fig 2a).

**Fig 2.**
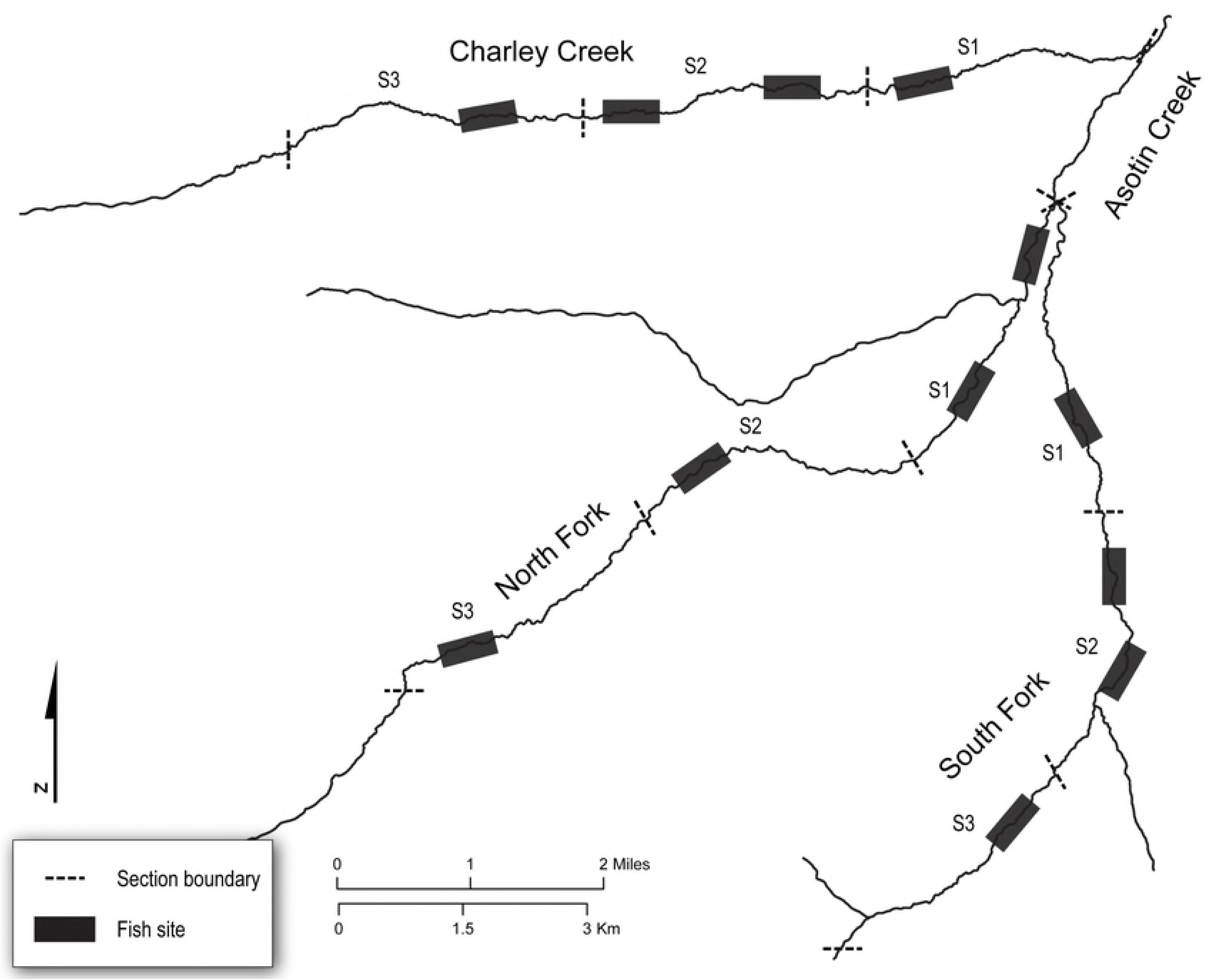
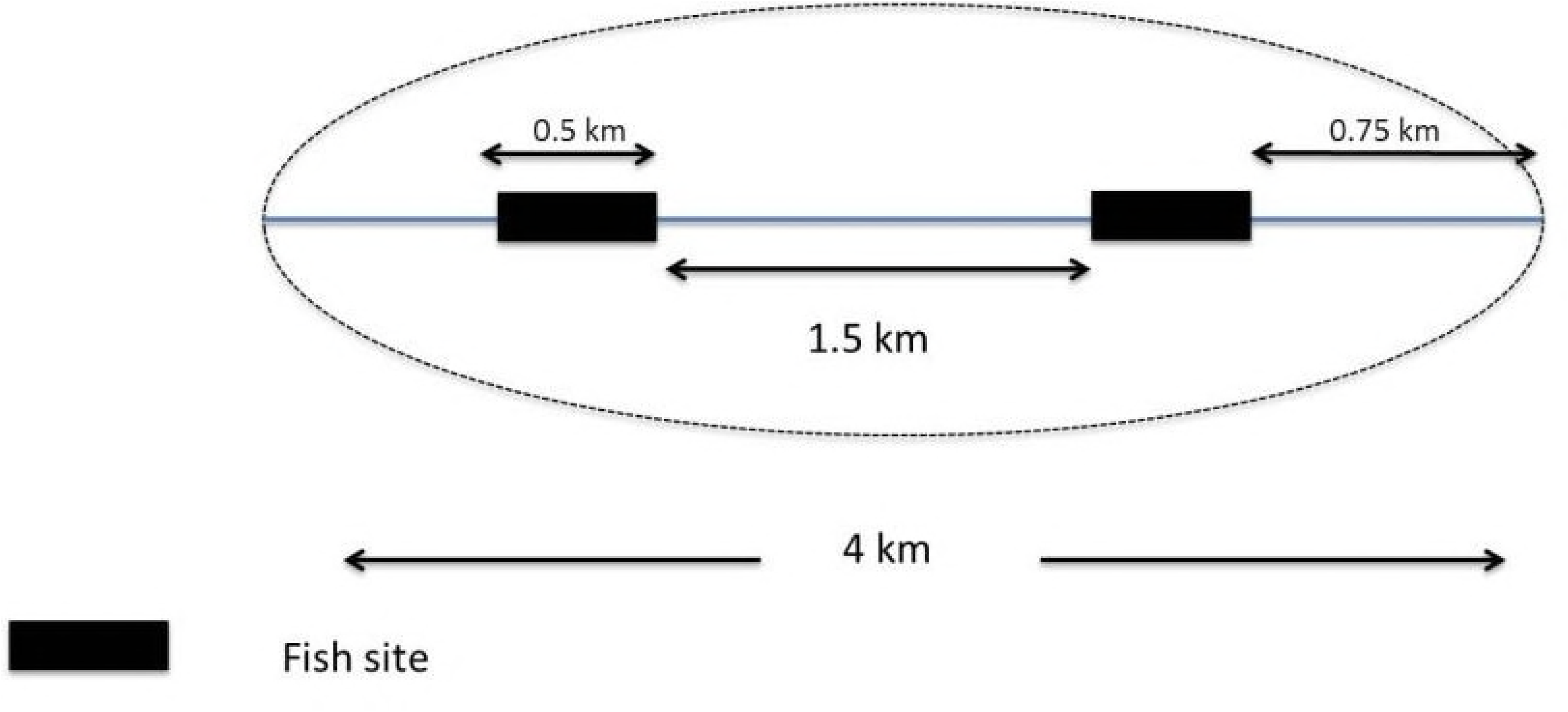
Layout of study design with three streams, three sections per stream, and six fish sample sites per section. a) Study design - each stream has three treatment units (4 km long *Sections*), and six fish sample sites (0.5 km long *Sites*). Timing and location of sections to be treated will depend on the study design implemented (see text for details). b) Survey design - each section has two fish sampling sites systematically located in each section 1.5 km apart.

Two sampling units (hereafter ‘fish sites’) for steelhead abundance, ~ 0.5 km long, were located systematically at 0.5 km and 2.5 km from the beginning of each experimental unit, resulting in all fish sites in a stream being 1.5 km apart (Fig 2b). We used a systematic sampling design to increase interspersion of the fish sites along the stream network to potentially detect longitudinal gradients and increase the likelihood that the survey locations were independent [27]. The mean and variance estimates used in the modeling simulation came from two day mark-recapture electrofishing surveys where all juvenile steelhead ≥ 70 mm are marked with passive integrated transponder tags and abundance estimated with the Chapman method [28]. Although we sampled juvenile steelhead abundance twice a year (summer and fall), we simplified the modeling by only using the summer abundance estimates for all simulations. We used density (fish/m^2^) as a measure of abundance for all simulations. We simulated different levels of sampling effort, but in general found that sampling effort had minimal and predictable effect on power. Increased sampling generally decreased confidence interval widths, regardless of experimental design. For the remainder of the paper, we only present the full sampling plan (two fish sites/section) to simplify the results and focus on the effect of the different designs.

### Simulation Approach

We simulated the Asotin IMW experiment by developing models that contained the various components of the IMW experiment. A simulation model was then developed for each of the four experimental designs by defining the terms (i.e., we defined “truth”) in the ANOVA model that would be used to analyze the data produced from these experiments. We identified stream, section within stream (written “section(stream)”), and fish site within section of stream (“fish-site(section x stream)”) as spatial factors and year, and the periods before or after the treatment, referred to as the *period,* as temporal factors (Table 1). We used historical data (1983-2006) and preliminary data from the Asotin IMW (2008-2009) on juvenile steelhead abundance in the study streams and in the mainstem Asotin creek to estimate the fixed effects (means) for the three streams and the variance components for all random effects (WDFW and S. Bennett, unpublished data). In all data sets, we observed that the variability of abundance measurements grew with larger mean abundances. The log transformation alleviated this problem and reduced the skewness of the measurements somewhat. We therefore applied all analysis and data simulation models to log(abundance) data.

**Table 1.**
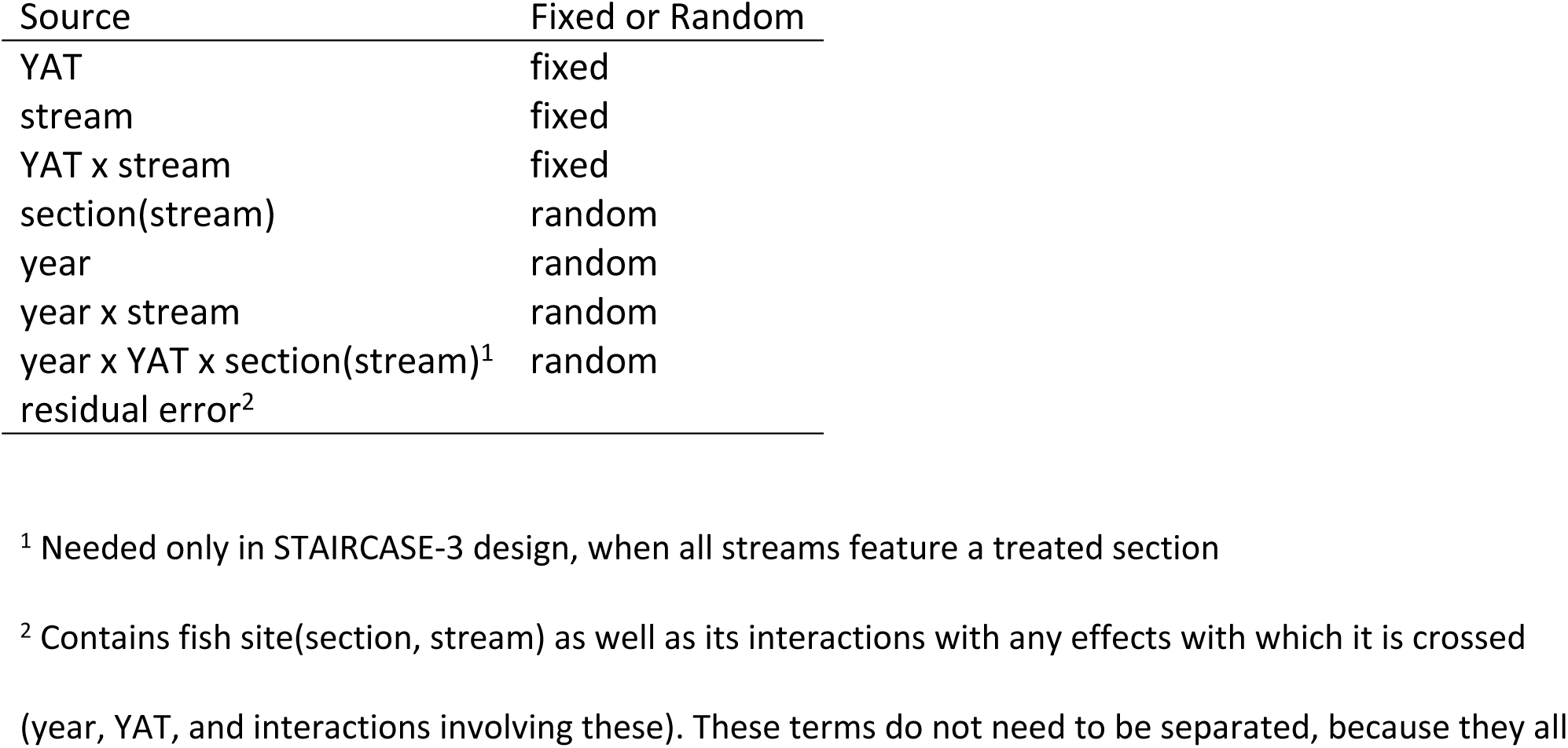
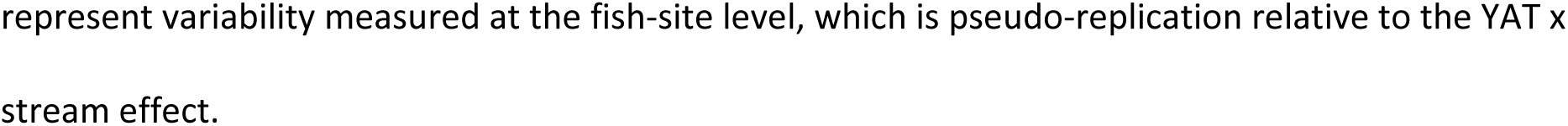
ANOVA model terms for simulation study of BACI and staircase designs.

| Source | Fixed or Random |
| --- | --- |
| YAT | fixed |
| stream | fixed |
| YAT x stream | fixed |
| section(stream) | random |
| year | random |
| year x stream | random |
| year x YAT x section(stream)<sup>1</sup> | random |
| residual error<sup>2</sup> |  |

<sup>1</sup> Needed only in STAIRCASE-3 design, when all streams feature a treated section
<sup>2</sup> Contains fish site(section, stream) as well as its interactions with any effects with which it is crossed (year, YAT, and interactions involving these). These terms do not need to be separated, because they all
represent variability measured at the fish-site level, which is pseudo-replication relative to the YAT x stream effect.

We used the historical data to estimate the mean log(abundance) in each stream and the corresponding *year*, *year x stream*, and *section(stream)* variance components (i.e., temporal variance; Table 2). We used our preliminary IMW fish data to estimate the *fish site(section, stream)* variance component. For the variance components, we additionally computed 95% Wald Confidence intervals around each estimate (Milliken and Johnson 2009). We used the estimates and confidence limits to define three different variability scenarios because the variance components were estimated with uncertainty that was sometimes substantial. We assumed that the estimated variance represented the “expected” variability present, and the lower and upper limit of the confidence intervals represented the “best case” and “worst case” for variability respectively (Table 2). Where a variance component could not be directly estimated from one of the existing data sets, educated guesses were used to fill in these missing values, according to the principle that smaller-size spatial or temporal units generally have smaller variance components (more similar responses) than larger size units (Milliken and Johnson 2009). Based on this review of temporal and spatial variance, we estimated that the watershed’s mean log (abundance) across years was 1.3 (CV = 37%). The mean log(abundance/m^2^) for the study streams were 1.46 for Charley Creek, 1.31 for North Fork Asotin Creek, and 1.35 for South Fork Asotin Creek.

**Table 2.** Variance components for log(abundance) based on historic juvenile steelhead abundance estimates from Washington Department of Fish and Wildlife (1986-2004) and preliminary sampling of Asotin IMW (2008-2009) for the different factors of the analysis. Expected is the variances calculated from each factor; Best case is the lower 95% confidence interval estimate of the variance; Worst case is the upper 95% confidence interval estimate of the variance.

| Factor | Expected | Best Case | Worst Case |
| --- | --- | --- | --- |
| year | 0.195 | 0.009 | 0.632 |
| section(stream) | 0.026 | 0.015 | 0.056 |
| fish site(section,stream) | 0.003 | 0.001 | 0.013 |
| year x stream | 0.006 | 0.002 | 0.560 |
| year x section(stream)<sup>12</sup> | 0.003 | 0.001 | 0.028 |
| year x fish site(section,stream)<sup>34</sup> | 0.002 | 0.000 | 0.006 |
| random error | 0.037 | 0.024 | 0.078 |

<sup>1</sup>Assumed to be $\frac{1}{2}$ x year x stream because section is subunit of stream. This effect could not be directly estimated from historical data. However, since it is both a spatial subunit of Year\*Stream and a temporal subunit of section(stream), we set it to $\frac{1}{2}$ of whichever of these two variance estimates was smaller, because subunits tend to have smaller variance components than the units within which they are nested.
<sup>2</sup> Assumed to be $\frac{1}{2}$ x section because changes in differences between sections across years are likely to be smaller than the original differences.
<sup>3</sup> Assumed to be $\frac{1}{2}$ x year x section because fish site is subunit of section. Term is a spatial subunit of year x section(stream) and a temporal subunit of site(section, stream), so entries were set to $\frac{1}{2}$ of the smaller of these two variances component estimates.
<sup>4</sup>Assumed to be $\frac{1}{2}$ x fish site because *changes* in differences between fish sites across years are likely to be smaller than the original differences.

We also checked our models for serial correlation across years on the measurements made within stream *sections* using an autoregressive process [AR(1); 29]. We found a statistically significant autocorrelation coefficient (r=0.42, 95% confidence interval 0.10-0.74) that we could then build into our simulations (see below).

### Simulation Modeling

We simulated four experimental designs based on the layout of the Asotin Creek lMW study defined above (Fig 2). First, we simulated an aBACI design in which one section was restored with LWD at the start of Year 7, and the eight other sections were controls (BACI-1; Fig 3). Second, we simulated an aBACI design where three sections were restored simultaneously with LWD at the start of Year 7, and the remaining six sections were maintained as controls (BACI-3; Fig 3). Third, we simulated a staircase experimental design where three sections of one stream were treated sequentially in time in a staircase fashion (STAIRCASE-1). During the first three years of the STAIRCASE-1 simulation all stream sections were left untreated, then, prior to measurements in each of Years 4, 7, and 10, one stream section was restored (Fig 3). Fourth, we simulated a staircase experimental design where the only difference with the STAIRCASE-1 simulation was one section from each stream was selected for treatment (STAIRCASE-3). The STAIRCASE-3 design has the advantage that each stream serves as its own control, so that the same experiment is essentially replicated three times in different years and streams.

**Fig 3.**
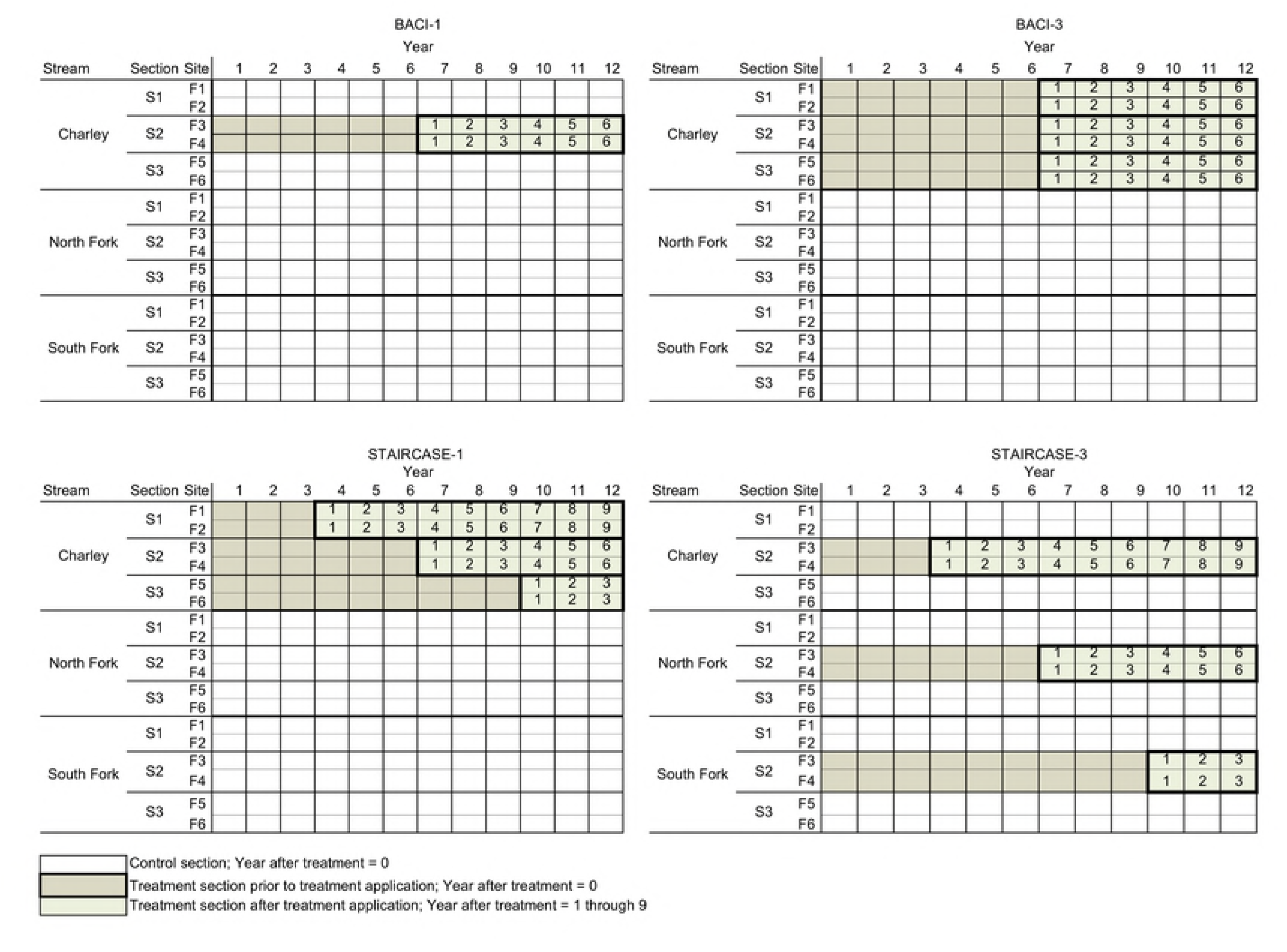
Schematic of four experimental designs and sample plans (i.e., all fish sites sampled). BACI-1 = one treatment section, BACI-3 = three simultaneous treatment sections in one stream, STAIRCASE-1 = three sequential treatment sections in one stream, and STAIRCASE-3 = three sequential treatment sections, one in each stream.

We simulated a range of period effects to develop power curves for the different variance assumptions. Among these factors, the *stream* was considered a fixed effect. The factors *year*, *section(stream)* and *fish-site(section x stream)* are considered as random effects factors. Although *section(stream)* were chosen deliberately and are not meant to be representative of other *section(stream)*, they act as random effects by virtue of having treatments randomly assigned to some number of them [30]. The ANOVA models given by Underwood (1992) for a variety of sampling designs are used with aBACI. The test for a restoration effect is obtained through the *treatment* x *period* interaction, where *period* is either 0 or 1 depending on whether the time of measurement is before or after the treatment application, respectively.

With a staircase design, the definition of *period* is more complicated than for aBACI design. A staircase design has *periods* for which there is a sliding definition of both “before” and “after,” and a given year may occur both before one treatment and after another. Also, control units serve as controls for all restored units, so that there is no clear way to match control units with treated units. This can make construction of a *treatment x period* interaction difficult. To address this, we matched the measurements on the restored units with respect to the time since the restoration was applied in their respective units. We define a fixed effect factor, *years after treatment* (YAT), which represents the number of years since restoration of a specific section. In our example, the Year 4 measurements on the earliest-restored section should be comparable to the Year 7 measurements on second-restored section and the Year 10 measurements on the last-restored section, because all of these measurements take place 1 year after restoration. Each of these measurements get YAT=1. Measurements taken on these sections in subsequent years get YAT=2, 3, … until the end of the study is reached. Notice that this limit is different for different restored sections because of the staggering of the establishment on different sections. Finally, all measurements prior to restoration being applied are considered to have YAT=0.

The ANOVA models used to analyze the simulation results have the standard assumptions that linear mixed models rely on, including normality of errors and random effects, additivity of the effects in the model, and independence of observations and all model effects [31, 32]. Our experiments contain five factors: one treatment factor (YAT) and four blocking factors (*year, stream, section(stream),* and *fish-site(section, stream)*). Note that *year* should not be treated as a longitudinal factor as is common in repeated-measures studies [33]. Responses among different spatial units are likely to be highly correlated with one another across years, as high and low fish abundance years affect all sections simultaneously (e.g. the effect of weather, or ocean effects on returns spawning fish). This invalidates the typical repeated-measures assumption that experimental units respond independently of one another (Loughin et al. 2007). Instead, time and space are crossed factors in the experimental design sense (Milliken and Johnson 2009), because the same series of years are observed on each spatial unit.

The terms contained in the ANOVA models are determined using the following standard conventions [30, 34]: all treatment and block factor main effects are included in the model; interactions among all crossed blocking factors are included in the model; and interactions between treatment and blocking factors are added discretionarily (Table 1). To keep our analyses as simple as possible and to give all study plans a common basis for comparison, we add no additional discretionary terms. Since YAT is assigned to the spatio-temporal unit defined by the crossing of year and *section(stream)*, the *year x section(stream)* effect becomes nested within *YAT*, forming the equivalent term *year x YAT x section(stream)* (Table 1). All analyses of the ANOVA model for each study plan were performed using PROC MIXED in SAS Version 9.3 (SAS Institute, Cary, NC). All SAS code is available from T. Loughin.

The model we used to generate data for the spatio-temporal structure of the IMW experiment was an additive normal linear model using terms based upon the analysis model described above. *Stream* and *YAT* effects were taken to be fixed; *year, section, and site* effects were taken to be random, and any interactions with random effects were taken to be random. Specifically, *year x section*, and *year x site* random effects were added.

Once the statistical model for the Asotin Creek IMW was defined, we simulated pseudo-random data to represent the potential measurement at each of 18 fish sites across 12 years in the Asotin IMW study area (hereafter the study area). Random effects were generated independently from zero-mean normal distributions according to their respective variance components, and individual stream (i.e., Charley, North Fork and South Fork) means were added in. A single simulation of an experimental design consisted of 216 potential measurements (2 fish sites/section x 3 sections/stream x 3 streams x 12 years = 216). We simulated 1000 model-watersheds for each of the 12 combinations of response variable, level of variability, and autocorrelation (see descriptions below). The number of simulations was chosen to allow Type I error rates of analyses conducted at the 5% level to be estimated to less than ±1.4% error with 95% certainty. Power estimates similarly can be estimated to at worst ±3.1% error with 95% certainty.

For each of the three variance scenarios, we used the simulation model to generate 1000 sets of log(abundance/m^2^) measurements that could be made if every fish site was sampled and no restoration was applied. We then reconfigured the potential responses according to each study plan as follows. We added treatment effects to the selected potential response values for all observations taken on restored sections in accordance with the experimental design being considered. Our first goal was to detect a 25% increase in mean abundance of juvenile steelhead in our treatment section(s), relative to our controls (within streams and between streams). Thus, log(1.25) was added to all treatment sections after restoration. However, to better understand the ability of each experimental design to detect a variety of treatment effects we varied the treatment effect from a 5% - 40% increase in juvenile steelhead abundance and developed power curves. This was intended to allow a more detailed comparison of the designs, specifically addressing the concern that multiple treatments applied in different sections of the same stream may synergize to generate a larger treatment effect in each restored section than would be observed by restoring only one section of a stream. For example, the hydraulics created by adding structures could result in downwelling that could increase hyporheic exchange. At the scale of a single structure this effect is likely to be minimal, but the scale of several hundred structures (multiple treated sections) could create a variability in water temperatures that could benefit juvenile steelhead [35]. By comparing the power curves for the four designs, we could observe how much synergy would need to take place to favor the single stream treatment designs (BACI-1, BACI-3, and STAIRCASE-1) over the three-stream treatment design (STAIRCASE-3).

We initially considered two possible patterns for treatment effects: “sudden” and “ramped”. For the sudden pattern, we assumed that the full treatment effect was observed in the first measurement subsequent to restoration and that this effect persisted through the remainder of the study. However, the addition of LWD to a stream may result in only incremental changes in the first few years after restoration, especially if spring flows are below average because high flows are necessary to produce the expected geomorphic response to alter fish habitat. Thus, for the ramped effect, we assumed that 60% of the full effect was realized in the first year, 90% in the second year, and 100% from the third year onwards. We found that the pattern of treatment had limited influence on the performance of the different study plans so we just present the results of the ramped treatment pattern because we feel it is a more realistic response. For simplicity, no specific *stream x treatment* interaction was created, because three of the experimental designs considered placed all treatments in one stream.

We designed linear contrasts to estimate a possible treatment effect. Generally, we designed contrasts to look for specific patterns in the treatment means. We chose basic contrasts that amounted to estimating mean effect of treatment across all YAT (compatible with seeking a “sudden” treatment effect). The model implicitly adjusted out any spatio-temporal effects from this estimate.

We used this estimate and its corresponding model-based standard error estimate to compute a 95% t-based confidence interval for the true treatment effect. We considered the treatment effect to be detected if the entire confidence interval was above 0 (i.e., the null hypothesis of no treatment effect would be rejected by a 2-sided t-test at the 0.05 significance level). We computed the proportion of simulations in which the treatment effect was detected at the given cutoff of p=0.05 for each experimental design. We also computed the mean length of the confidence intervals, because we are also interested in whether ability to detect a response was because of the precision or the size of the effect.

To estimate the influence of variance on the power and confidence interval for each experimental design, we used all three levels of variability (expected, best case, and worst case) calculated from our analysis of historic and trial data (Table 2). Each simulation contained the potential for serial autocorrelation because annual correlations were observed to exist in the abundance of juvenile steelhead in the different streams. We therefore simulated annual means (*year* effects) using autocorrelation of either 0 or 0.5.

## Results

Under the best-case variability, all designs had 100% power to detect a 25% increase in juvenile fish abundance. Power remained high under the expected variability as all designs had at least 94% power to detect the simulated 25% increase in abundance (Table 3). The power to detect a 25% increase in fish abundance under the worst-case variability was almost twice as high in the STAIRCASE-3 design (77%) compared to the BACI-1 (41%) and STAIRCASE-1 designs (33%). The BACI-3 design had the lowest power to detect a 25% increase in fish abundance at only 8%. All designs detected the simulated 25% increase in abundance accurately under expected and worst-case variability (Fig 4a and b). Confidence interval lengths were similar for all designs under expected variance (Fig 4 a); however, BACI-3 had larger confidence interval lengths under both expected and worst-case variability and only STAIRCASE-3 had confidence intervals that did not cross 0 under worst-case variability (Fig 4b).

**Fig 4.**
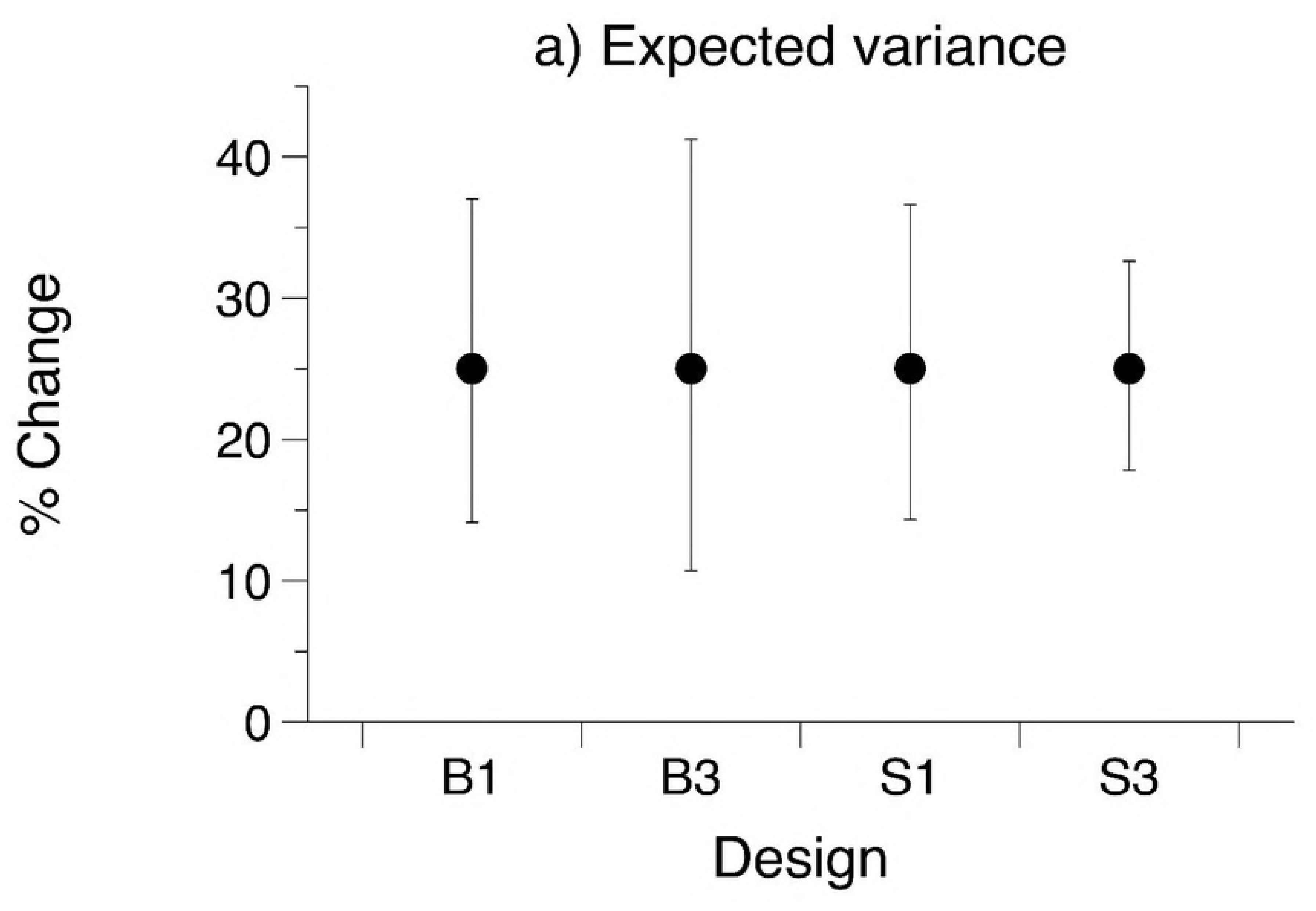
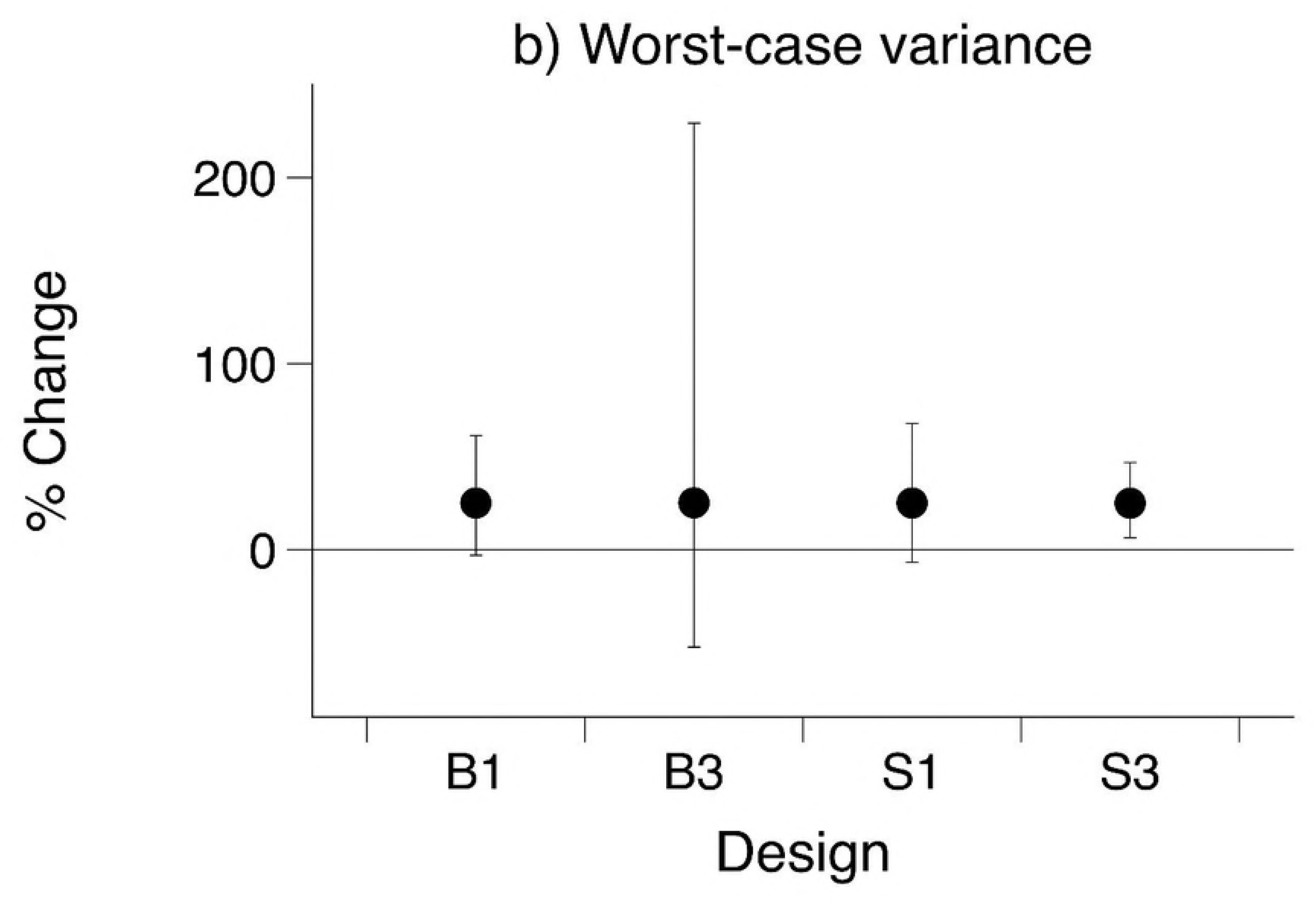
Estimated percent change in juvenile steelhead abundance and confidence intervals from a simulated 25% change in abundance for four experimental designs under a) expected and b) worst-case variability. BACI-1 = one treatment section, BACI-3 = three simultaneous treatment sections in one stream, STAIRCASE-1 = three sequential treatment sections in one stream, and STAIRCASE-3 = three sequential treatment sections, one in each stream.

**Table 3.** Summary of the power, lower and upper 95% confidence intervals, and power to detect a 25% increase in juvenile fish abundance under expected and worst-case variance for four different experimental designs.

| Variance | Design | 95% Confidence Intervals |  |  | Power |
| --- | --- | --- | --- | --- | --- |
|  |  | % Change | Low | Up |  |
| Expected | BACI-1 | 25.0 | 14.1 | 37.0 | 1.00 |
|  | BACI-3 | 25.0 | 10.7 | 41.2 | 0.94 |
|  | STAIRCASE-1 | 25.0 | 14.4 | 36.7 | 1.00 |
|  | STAIRCASE-3 | 25.0 | 17.8 | 32.7 | 0.00 |
| Worst | BACI-1 | 25.1 | -3.0 | 61.3 | 0.41 |
|  | BACI-3 | 25.2 | -52.4 | 229.3 | 0.08 |
|  | STAIRCASE-1 | 25.1 | -6.8 | 67.8 | 0.33 |
|  | STAIRCASE-3 | 24.9 | 6.3 | 46.8 | 0.77 |

We generated power curves to assess the ability of each design to detect smaller changes in abundance (Fig 5). STAIRCASE-3 design consistently had higher power to detect a range of abundance increases, BACI-3 consistently had the lowest power, and BACI-1 and STAIRCASE-1 had similar power under expected and worst-case variance. For example, a 10% increase in abundance with expected variance resulted in power from a high of 88% power (STAIRCASE-3) to a low of 34% power (BACI-3; Fig 5a). The power to detect a 10% increase in abundance with worst-case variance resulted in power ranging from a high of 21% (STAIRCASE-3) to a low of 5% (BACI-3; Fig 5b).

**Fig 5.**
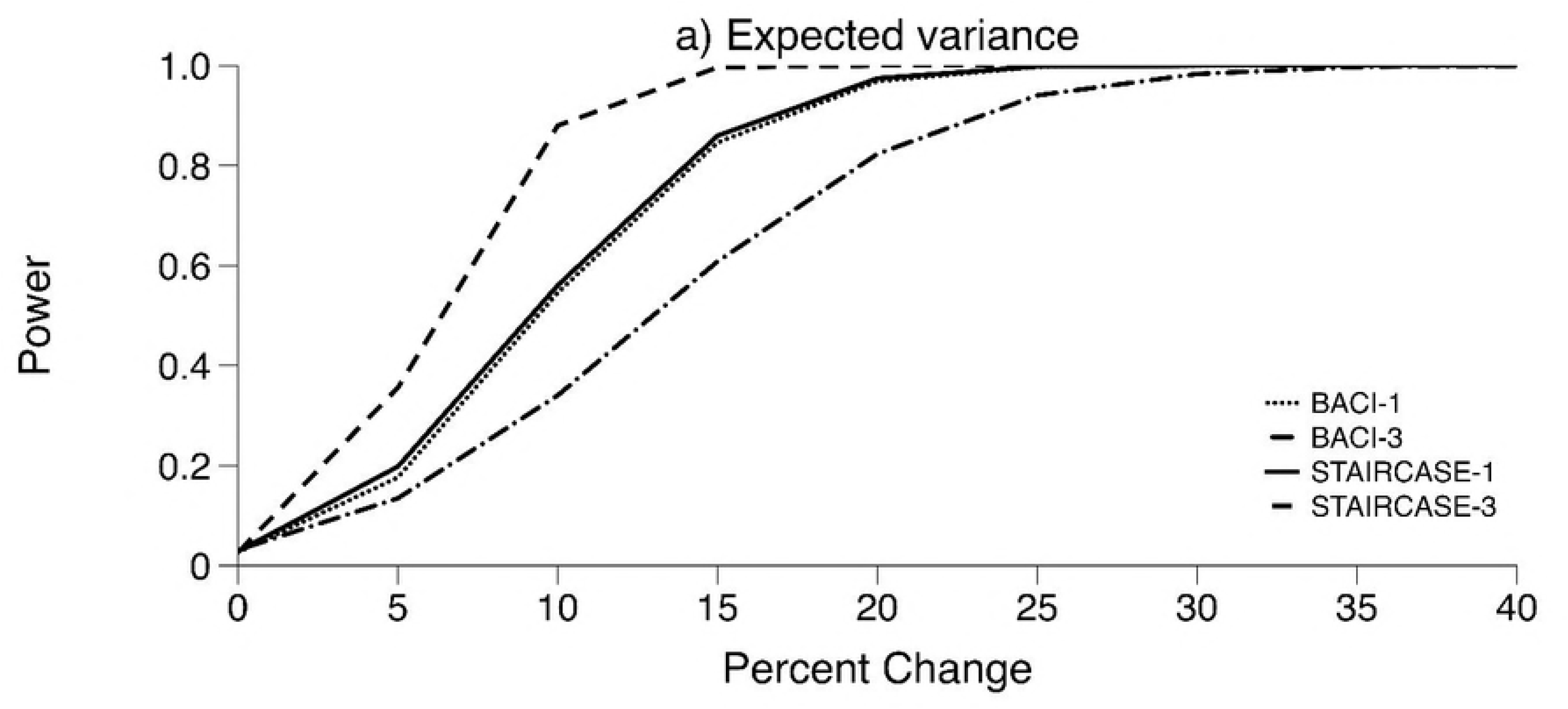
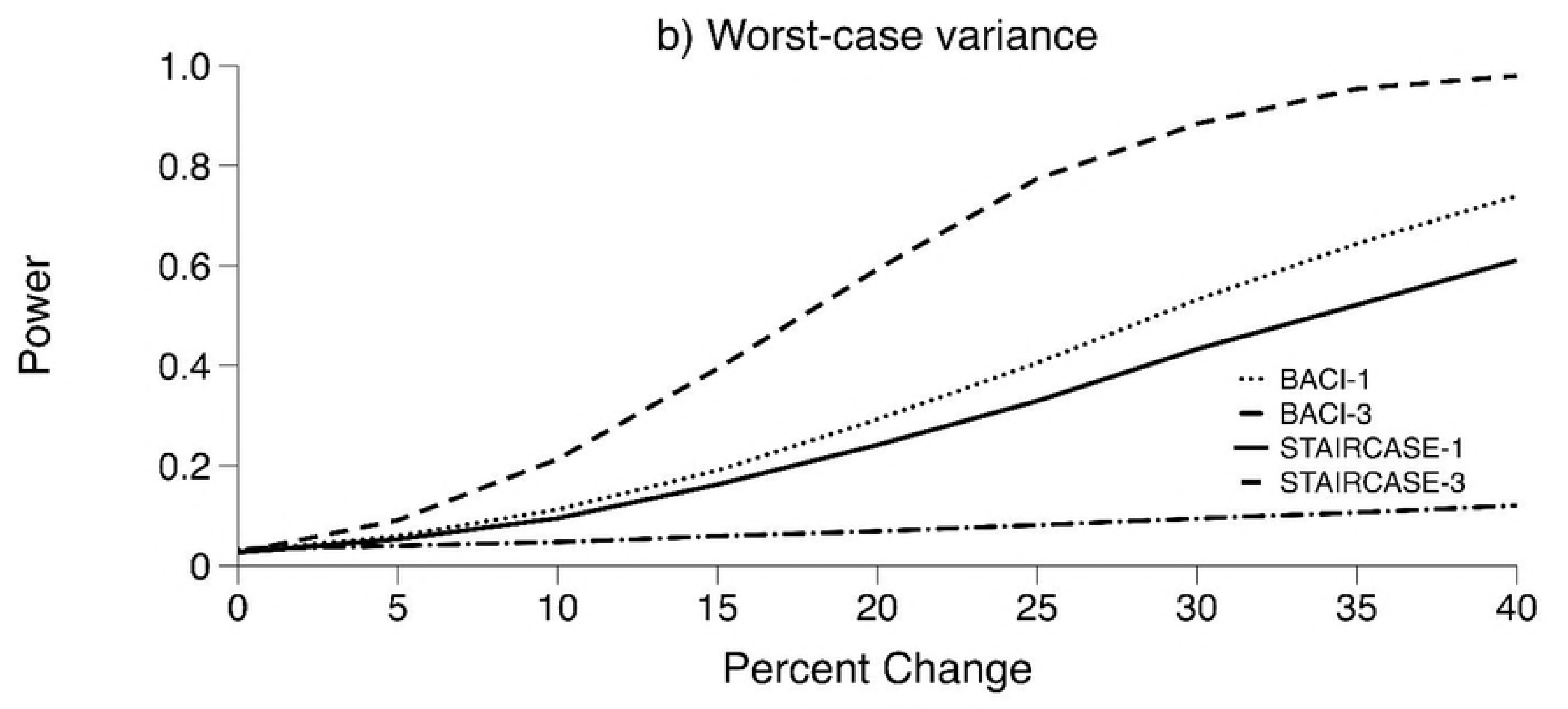
Power curves for four experimental designs with varying sizes of treatment effect under a) expected variability and b) worst-case variability. BACI-1 = one treatment section, BACI-3 = three simultaneous treatment sections in one stream, STAIRCASE-1 = three sequential treatment sections in one stream, and STAIRCASE-3 = three sequential treatment sections, one in each stream.

The BACI-3 and STAIRCASE-1 designs have the potential to have similar power to detect changes in abundance if treating three sections in one stream causes a greater overall response than treating three sections in three different streams (i.e., synergy of continuous treatments). Under expected variability, the synergy would need to result in a 60-150% greater increase in abundance to make the power curves similar for BACI-3, STAIRCASE-1, and STAIRCASE-3. However, the synergy would have to be greater under worst-case variability because the confidence intervals are independent of the size of a treatment effect and only STAIRCASE-3 has confidence intervals that do not cross 0. Therefore, in order for the BACI-3 and STAIRCASE-1 design to have similar power to the STAIRCASE-3 design there would need to be considerable synergy—generally about 300% the increase in abundance (Fig 5b).

## Discussion

We developed four experimental designs and used field estimates of the mean and variance of fish abundance to develop a simulation model where we assessed the power of each design to detect a response to stream restoration by defining “truth” to generate data that was then analyzed by the appropriate statistical models. We demonstrated that the staircase design could have significantly more power to detect treatment responses than aBACI designs, especially when variance levels are high or the size of the effect is small. An important feature of staircase designs is that they are more robust against unusual circumstances in a given year that might cause atypical responses in one or more treatment or control groups. For example, in experiments like ours, the timing or magnitude of spring floods can alter the treatment effect dramatically. These year-by-treatment effects can confound experiments in which there is only a single time of treatment application (Walters et al. 1988). Walters et al. (1988) and Loughin et al. (2007) demonstrated that staggering the start of long-term experiments could guard against such problems. Despite this advantage, few examples exist in the literature where staircase designs have been applied to large-scale experiments.

Another benefit of the staircase design over a whole stream manipulation is that logistically, application of restoration in sections staggered over time is more feasible than over entire stream in a single year. Regardless, the whole stream manipulation (BACI-3) resulted in very low power to detect an effect under most scenarios. Furthermore, we found that the BACI-3 design was less powerful than the BACI-1 design, which is even more logistically feasible than the staircase designs. Although not as powerful as the STAIRCASE-1 and STAIRCASE-3 designs, the BACI-1, could be a reasonable approach if the effect is large and the variability is moderate.

Criticisms that the benefits of stream restoration are potentially misleading or hard to detect stems from the localized evaluation of restoration effectiveness (i.e. the structure or small reach scale; Thompson 2005, Katz et al. 2007). The concerns are that small-scale restoration and evaluation might result in fish being “attracted” to these small changes with no real population benefit, or that the size of the effects are too small to detect. To overcome such issues, assertions that the entire stream (watershed) needs to be restored in order to detect a true fish response [11, 12]. Indeed, several studies evaluating restoration benefits follow this approach (Bennett et al. 2016). Thus, the low power of the BACI-3 is somewhat unintuitive given this assertion.

The low power of the BACI-3 occurs in part because we have assumed there exists a *stream by year* interaction as supported by our analysis of existing data. That is, the fish populations in different streams within our watershed do not change identically between years, even though they all experience similar out-of-watershed effects (e.g. the effects of the ocean, fishing, and Columbia and Snake hydropower system), and localized climate effects. While using multiple streams can greatly reduce the noise caused by these factors, noise created by asynchrony between streams is likely to always exist. Keeping that extraneous noise out of the treatment comparisons is one of the key goals of an experimental design. In much the same way as blocking is used in typical experimental designs to improve precision of treatment comparisons, maintaining both control and treated sections in streams improves the power for our IMW designs (Underwood 1994). Therefore, if all the treatments are applied to one stream and no controls are maintained within the stream, the comparisons between treatments must be made between streams as well, rather than between sections within the same stream. The variance caused by *stream by year* interaction confounds with the treatment effects, making the treatment effects more difficult to detect.

Similarly, STAIRCASE-3 performs better than STAIRCASE-1 because STAIRCASE-1 loses an internal control once all the reaches are treated. The temporal variability of sections is the most important component of the error term for testing and forming confidence intervals for treatment effects. Increases in temporal variance results in increases in the variability of the treatment effects, especially those associated with times more than 6 years after treatment. The advantages of multiple treated sections are diminished by the disadvantages of increased difficulty in separating treatment effects from inherent variability. The STAIRCASE-3 design overcomes this issue by having treated sections spread among three streams, with control sections in the same streams. This creates a situation akin to blocking in that treatment comparisons against controls are made within stream rather than between streams, and therefore incur less variability in estimating effects, resulting in the increase in power and shorter confidence intervals of the STAIRCASE-3 design.

The analyses include terms that account for any variability that occurs on a larger scale, and hence this variability does not affect the designs’ relative powers or confidence interval lengths. To check this assumption, we inflated individual variance components and reran some simulations. We found no change in power when we inflated variance components that correspond to year interactions with units larger than *section*. We also assumed that a 4km section of stream represents a “population” or is functionally independent of other sections. This assumption allows for greater replication in treatments and controls for design considerations. We can therefore evaluate the expected response of a stream from one section, while keeping internal controls within a stream to evaluate the *stream by year* interaction. In the STAIRCASE-3 design, we are essentially replicating an aBACI three times. We assumed that the addition of LWD would create local increases in habitat complexity that would be unlikely to produce changes in habitat complexity in upstream or downstream sections. If section-to-section movement of fish remained low as determined from previous data, then we can expect that the treatment effects on abundance will be mostly independent as well. As we will discuss below, we could test this hypothesis more stringently as the experiment proceeds.

If we believe treatments applied to different sections of the same stream do synergize to create a broader, more favorable environment for fish, then the application of multiple treatments to sections of the same stream has the potential to create a larger effect that is easier to detect than effects caused by other designs. However, we have shown that the increases due to synergism would need to be considerable for the BACI-3 and STAIRCASE-1 designs to match the power of the STAIRCASE-3 design.

A similar important consideration that we did not pursue in the study is whether the interpretation of results is affected if the assumption of treatment independence is violated. The results of our simulation could be confounded if the degree of movement of fish between reaches is not well known and more mixing occurs between treatment and control reaches [27]. Alternatively, the manipulation may affect multiple stream sections, which might be particularly true when a treatment occurs upstream of the control sections. Underwood (1994) suggest that when the scale of the impact is unknown a hierarchical experimental design can be useful in identify the scale of the impact and response, and protect against losing controls when finer scale controls are no longer independent (i.e. replicates are pooled to watershed scale comparisons). For example, in Bridge Creek, Bouwes, Weber (36) evaluated the impacts of beaver dams on steelhead production by comparing reaches installed with beaver dam analogs (BDAs) to control reaches within a treatment watershed. A control watershed was also monitored for the same steelhead responses. Beavers quickly built dams on and between BDAs, and then moved into the control reaches and built more dams, compromising the ability to observe differences between treatment and control reaches. Therefore, all reaches were pooled into the treatment watershed and compared to the control watershed, revealing large benefits of beaver dams to the steelhead population. Future simulations could compare the ability to detect a stream level response in the event that restoration impacts are far reaching, beyond the section treated. The multiple designs could be compared across a range of assumptions to determine the trade-off and robustness of different designs.

We used abundance of juvenile steelhead for our simulations. However, many experiments focused on testing the effectiveness of restoration are interested in metrics that are tied to fitness, such as growth, survival, productivity (i.e., smolts/spawner), and production (g/area/time); all of which are likely to be far more variable because of the number of individuals that have to be tagged and recaptured to produce reliable metrics [36]. The determination of the size of the treatment section should evaluate the trade-offs of tagging enough fish, while maintaining enough independence where movement between sections has little effect on the different metric means and variability, to maximize experimental replicates. As this study illuminates, these kind of analyses could be used to challenge the assumption that larger-scale treatments make it easier to detect restoration effectiveness.

This study reveals that staircase designs can be powerful alternative to BACI designs, especially under worst-case variability and small effect sizes. However, the BACI-1 can also be a powerful design in these circumstances. Further simulations could help elucidate what happens when certain assumptions are violated, such as the degree that the loss of independence has on detecting effects. Such analyses are highlighted in classical adaptive management examples [37, 38]. However, the planning of similar restoration experiments may not have the level of data we had to carry out analyses. Given our results, a prudent approach to maximizing learning while minimizing costs, would be to collect pre-treatment data and evaluate treatment section length by assessing the trade-offs of fish sample size and independence as described above. Then treat this section size, and continue data collection to evaluate the agreement of assumptions. If the effect size appears to be large (i.e. large effect or low CV) while maintaining independence, the BACI-1 might be the appropriate approach as it requires the least expense in terms of evaluating restoration effectiveness. If the effect size (i.e. small effect or high CV) appears to be small, and independence is maintained, then the STAIRCASE-3 design is most likely to be most appropriate. If fish movement or treatment effects extend into the other sections, then the STAIRCASE-1 design is likely to be the most powerful. However, if the goal is to maximize both learning and recovery efforts, then STAIRCASE-3 or STAIRCASE-1 will be a clear choice over BACI-1 and BACI-3. As promoted in adaptive management, these goals and the decision making process should be explicitly stated prior to implementation of the experiment to avoid making this effort a trial and error process. In this fashion, application of the experiment and intensive monitoring can be an alternative to extensive modeling and evaluation of pre-treatment data as in classic examples of adaptive management (Bouwes et al. 2016a).

The mixed models we developed for our study plans could be easily extended to other experimental and sampling designs. Indeed, the mixed-model paradigm is quite flexible and could be adapted to other study designs by careful identification of the units on which fixed and random effects are measured [30, 39, 40]. We used historical data from the same watershed to inform the model for our simulated watersheds, so the data produced by the simulation should be reasonably representative of the potential measurements on this watershed (at least, as much as any empirical model can represent such a complex hydrological and ecological environment).

Defining year after treatment (*YAT*) allowed us to be flexible in creating treatment-effect patterns and to create linear contrasts [direct comparisons of appropriately selected means; 31] to estimate these different patterns. For example, if we want to estimate the mean response after treatment (a “press” effect sensu Underwood 1992), we can write a contrast that places equal weight on each YAT level above 0. If we want to estimate the effect only in the first year, or second year (etc.) after treatment (a “pulse” effect sensu Underwood 1992), we can write a contrast that places equal weight on the involved YAT level and zero elsewhere. We can define patterns of linear increase or decrease or many other patterns that match what we might expect our treatment effects to follow. Focusing on defining effects in this way also allows us to obtain numerical estimates of treatment effects (e.g., mean differences) and corresponding confidence intervals, rather than to rely on hypotheses tests that are not nearly as informative [41–43].

The principles that drive the comparisons among the power of different designs do not depend on the actual values of the variance components, but rather on their relative sizes. Fundamentally, treatments are applied to sections and subsequently measured in different years.

## Conclusion

Large investments have been made in restoring stream habitat with the explicit assumption that the restoration is increasing salmon and steelhead productivity [44]. In order to determine the effectiveness of restoration, well-designed experiments using restoration as the treatment are required. Traditional BACI designs have improved since their first development and are now commonly used in restoration experiments and assessments of environmental impacts [45]. Despite the wide-spread use of BACI designs, there are still debates in the literature about their proper use. We have presented a staircase design that has several statistical and logistical advantages over BACI designs. BACI designs that explicitly treat a whole watershed may have low power because of a loss of internal controls. In situations where variability is high, staircase designs that apply restoration to multiple streams have considerably higher power to detect changes in treatment responses. The potential hierarchical nature of the staircase designs also make them well suited for trying to understand systems where the scale of the treatment response is unclear and maximizing learning about the study subjects is critical in an adaptive management framework. Studies measuring the response of salmon and steelhead populations to restoration are ideal for using staircase designs. Despite the increased complexity in the design, implementation, and analysis of such experiments, our results suggest that there are significant benefits to using these designs, especially toward the primary goal of IMWs to answer the question, “does restoration work?” We strongly recommend that researchers utilize staircase designs in future projects but implement them in an adaptive management framework to maximize learning [36, 44].

## Acknowledgements

We thank Pete McHugh for reviewing early versions of this manuscript, Doris Bennett for editing, and Nick Weber and Chalese Hafen for help creating graphics.

**Author contributions** Nick Bouwes conceptualized the research and questions this manuscript addressed. Tom Loughin developed the methodology and conducted the simulations, developed the statistical models, formal analysis, and prepared the initial manuscript draft. Stephen Bennett developed the experimental and survey designs, conducted the initial investigations, visualization material, and led the writing, review, and editing of the final manuscript.

## References

1. Likens GE, Bormann FH, Johnson NM, Fisher DW, Pierce RS. Effects of forest cutting and herbicide treatment on nutrient budgets in the Hubbard Brook Watershed-Ecosystem. Ecol Monogr. 1970;40:23–47.

2. Carpenter SR. Large-scale perturbations: opportunities for innovation. Ecology. 1990;71(6):2038–43.

3. Hartman GF, Scrivener JC, Miles MJ. Impacts of logging in Carnation Creek, a high-energy coastal stream in British Columbia, and their implications for restoring fish habitat. Canadian Journal of Fisheries and Aquatic Sciences. 1996;53(1):237–51.

4. Carpenter SR, Chisholm SW, Krebs CJ, Schindler DW, Wright RF. Ecosystem experiments. Science. 1995;269(5222):324–7.

5. Carpenter SR, Frost TM, Heisey D, Kratz TK. Randomized intervention analysis and the interpretation of whole-ecosystem experiments. Ecology. 1989;70(4):1142–52.

6. Bennett S, Pess G, Bouwes N, Roni P, Bilby RE, Gallagher S, et al. Progress and Challenges of Testing the Effectiveness of Stream Restoration in the Pacific Northwest Using Intensively Monitored Watersheds. Fisheries. 2016;41(2):92–103. doi:10.1080/03632415.2015.1127805.

7. Katz SL, Barnas K, Hicks R, Cowen J, Jenkinson R. Freshwater habitat restoration actions in the Pacific Northwest: a decade’s investment in habitat improvement. Restor Ecol. 2007;15(3):494–505. doi:doi:10.1111/j.1526-100X.2007.00245.x.

8. Thompson DM. The history of the use and effectiveness on instream structures in the United States. In Ehlen, J Haneberg, WC, and RA Larson Eds Humans as Geologic Agents, Boulder, CO Geological Society of America Reviews in Engineering Geology. 2005;XVI: 35–50.

9. Kondolf GM, Anderson S, Lave R, Pagano L, Merenlender A, Bernhardt ES. Two decades of river restoration in California: what can we learn? Restor Ecol. 2007;15(3):516–23. doi:doi:10.1111/j.1526-100X.2007.00247.x.

10. Roni P, Hanson K, Beechie T. Global review of physical and biological effectiveness of stream habitat rehabilitation techniques. North American Journal of Fisheries Management. 2008;28:856–90.

11. Roni P, Pess G, Beechie T, Morley S. Estimating changes in Coho Salmon and Steelhead abundance from watershed restoration: how much restoration is needed to measurably increase smolt production? North American Journal of Fisheries Management. 2010;30(6):1469–84. doi:10.1577/m09-162.1. PubMed PMID: WOS:000286421000014.

12. Liermann M, Roni P. More sites or more years? Optimal study design for monitoring fish response to watershed restoration. North American Journal of Fisheries Management. 2008;28(3):935–43. PubMed PMID: ISI:000257780100030.

13. Green RH. Sampling design and statistical methods for environmental biologists. John Wiley and Sons, New York. 1979.

14. Smith EP, Orvos DR, Cairns JJ. Impact Assessment Using the Before-After-Controi-Impact (BACI) Model: Concerns and Comments. Canadian Jiournal of Fisheries and Aquatic Sciences. 1993;50:627–37.

15. Underwood AJ. Beyond BACI: experimental designs for detecting human environmental impacts on temporal variations in natural populations. Australian Journal of Marine and Freshwater Research. 1991;42:569–87.

16. Underwood AJ. On beyond BACI: sampling designs that reliably detect environmental disturbances. Ecological Applications. 1994;4(1):3–15.

17. Korman J, Higgins B. Evaluating habitat restoration on salmon abundance. Canadian Journal of Fisheries and Aquaitc Sciences. 1997;54:2058–67.

18. Johnson SL, Rodgers JD, Solazzi MF, Nickelson TE. Effects of an increase in large wood on abundance and survival of juvenile salmonids (Oncorhynchus spp.) in an Oregon coastal stream. Canadian Journal of Fisheries and Aquatic Sciences. 2005;62:412–24.

19. Walters CJ, Collie JS, Webb T. Experimental designs for estimating transient responses to management disturbances. Canadian Journal of Fisheries and Aquatic Sciences. 1988;45(3):530–8.

20. Loughin TM, Roediger MP, Milliken GA, Schmidt JP. On the analysis of long-term experiments. Journal of the Royal Statistical Society: Series A (Statistics in Society). 2007;170(1):29–42.

21. Jelks HL, Walsh SJ, Burkhead NM, Contreras-Balderas S, Díaz-Pardo E, Hendrickson DA, et al. Conservation status of imperiled North American freshwater and diadromous fishes. Fisheries. 2008;33(8):372–407.

22. Bennett S, Bouwes N. Southeast Washington Intensively Monitored Watershed Project: Selection Process and Proposed Experimental and Monitoring Design for Asotin Creek. State of Washington, Recreation and Conservation Office, Olympia, Washington. 2009.

23. Wheaton J, Bennett S, Bouwes B, Camp R. Asotin Creek Intensively Monitored Watershed: Restoration plan for Charley Creek, North Fork Asotin, and South Fork Asotin Creeks. DRAFT: April 7, 2012. Prepared for the State of Washington Recreation and Conservation Office, Olympia, WA Prepared by Eco Logical Research Ltd. 2012.

24. McIntosh BA, Sedell JR, Smith JE, Wissmar RC, Clarke SE, Reeves GH, et al. Management History of Eastside in Fish Habitat over 50 years 1935-1992. USDA Forest Service General Technical Report. 1994.

25. Fox M, Bolton S. A regional and geomorphic reference for quantities and volumes on instream wood in unmanaged forested basins of Washington State. North American Journal of Fisheries Management. 2007;27:342–59.

26. Camp RJ. Short-term effectiveness of high density large woody debris, a cheap and cheerful restoration action, in Asotin Creek. Master’s thesis. Utah State University, Logan, Utah. 2015.

27. Hurlbert SH. Pseudoreplication and the design of ecological field experiments. Ecological Monographs. 1984;54(2):187–211.

28. Seber GAF. A Review of Estimating Animal Abundance .2. Int Stat Rev. 1992;60(2):129–66. PubMed PMID: ISI:A1992JG51400002.

29. Littell RC, Stroup WW, Milliken GA, Wolfinger RD, Schabenberger O. SAS for Mixed Models, Second Edition. SAS Institute, Cary, NC. 2006.

30. Piepho HP, Büchse A, Emrich K. A hitchhiker’s guide to mixed models for randomized experiments. Journal Agronomy & Crop Science. 2003;189:310–22.

31. Milliken GA, Johnson DE. Analysis of Messy Data: Designed Experiments, Volume 1, Second Edition. CRC Press, Boca Raton, FL, USA. 2009.

32. Underwood AJ. Techniques of analysis of variance in experimental marine biology and ecology. Annual Review of Oceanography and Marine Biology. 1981;19:513–605.

33. Green RH. Application of repeated measures designs in environmental impact and monitoring studies. Australian Journal of Ecology. 1993;18:81–98.

34. Nelder JA. The Analysis of Randomized Experiments with Orthogonal Block Structure. I. Block structure and the Null Analysis of Variance. Proceedings of the Royal Society of London Series A, Mathematical and Physical Sciences. 1965;283:147–62.

35. Weber N, Bouwes N, Pollock MM, Volk C, Wheaton JM, Wathen G, et al. Alteration of stream temperature by natural and artificial beaver dams. PLoS One. 2017;12(5):e0176313. doi:10.1371/journal.pone.0176313. PubMed PMID: 28520714; PubMed Central PMCID: PMCPMC5435143.

36. Bouwes N, Weber N, Jordan CE, Saunders WC, Tattam IA, Volk C, et al. Ecosystem experiment reveals benefits of natural and simulated beaver dams to a threatened population of steelhead (Oncorhynchus mykiss). Scientific reports. 2016;6:28581. doi:10.1038/srep28581. PubMed PMID: 27373190; PubMed Central PMCID: PMCPMC4931505.

37. Walters C. Adaptive management of renewable resources. McMillan Publishing, New York. 1986.

38. Holling CS. Adaptive environmental assessment and management. Wiley, Chichester, U.K. 1978.

39. Brien CJ, Bailey RA. Multiple randomizations (with discussion). Journal of the Royal Statistical Society, Series B. 2006;68:571–609.

40. Brien CJ, Demetrio CBG. Formulating mixed models for experiments, including longitudinal experiments. Journal of Agricultural, Biological, and Environmental Statistics. 2009;14:253–80.

41. Matloff NS. Statistical hypothesis testing: problems and alternatives. Environ Entomol. 1991;20(5):1246–50.

42. Suter GW. Abuse of hypothesis testing statistics in ecological risk assessment. Human and Ecological Risk Assessment. 1996;2(2):331–47. PubMed PMID: WOS:A1996WC99100013.

43. Anderson DR, Burnham KP, Thompson WL. Null hypothesis testing: probelms, prevalance, and an alternative. Journal of Wildlife Management. 2000;64(4):912–23.

44. Rieman BE, Smith CL, Naiman RJ, Ruggerone GT, Wood CC, Huntly N, et al. A comprehensive approach for habitat restoration in the Columbia Basin. Fisheries. 2015;40(3):124–35. doi:10.1080/03632415.2015.1007205.

45. Smokorowskia KE, Randall RG. Cautions on using the Before-After-Control-Impact design in environmental effects monitoring programs. Facets. 2017;2(1):212–32. doi:10.1139/facets-2016-0058.

